# Dissecting the Roles of Kalirin-7/PSD95/GluN2B Interactions in Different Forms of Synaptic Plasticity

**DOI:** 10.1101/744508

**Authors:** Mason L. Yeh, Jessica R. Yasko, Eric S. Levine, Betty A. Eipper, Richard E. Mains

## Abstract

Kalirin-7 (Kal7) is a Rac1/RhoG GEF and multidomain scaffold localized to the postsynaptic density which plays an important role in synaptic plasticity. Behavioral and physiological phenotypes observed in the Kal7 knockout mouse are quite specific: genetics of breeding, growth, strength and coordination are normal; Kal7 knockout animals self-administer cocaine far more than normal mice, show exaggerated locomotor responses to cocaine, but lack changes in dendritic spine morphology seen in wildtype mice; Kal7 knockout mice have depressed surface expression of GluN2B receptor subunits and exhibit marked suppression of long-term potentiation and depression in hippocampus, cerebral cortex, and spinal cord; and Kal7 knockout mice have dramatically blunted perception of pain. To address the underlying cellular and molecular mechanisms which are deranged by loss of Kal7, we administered intracellular blocking peptides to acutely change Kal7 function at the synapse, to determine if plasticity deficits in Kal7^-/-^mice are the product of developmental processes since conception, or could be detected on a much shorter time scale. We found that specific disruption of the interactions of Kal7 with PSD-95 or GluN2B resulted in significant suppression of long-term potentiation and long-term depression. Biochemical approaches indicated that Kal7 interacted with PSD-95 at multiple sites within Kal7.

**Graphical Table of Contents:** The postsynaptic density is an integral player in receiving, interpreting and storing signals transmitted by presynaptic terminals. The correct molecular composition is crucial for successful expression of synaptic plasticity. Key components of the postsynaptic density include ligand-gated ion channels, structural and binding proteins, and multidomain scaffolding plus enzymatic proteins. These studies address whether the multiple components of the synaptic density bind together in a static or slowly adapting molecular complex, or whether critical interactions are fluid on a minute-to-minute basis.

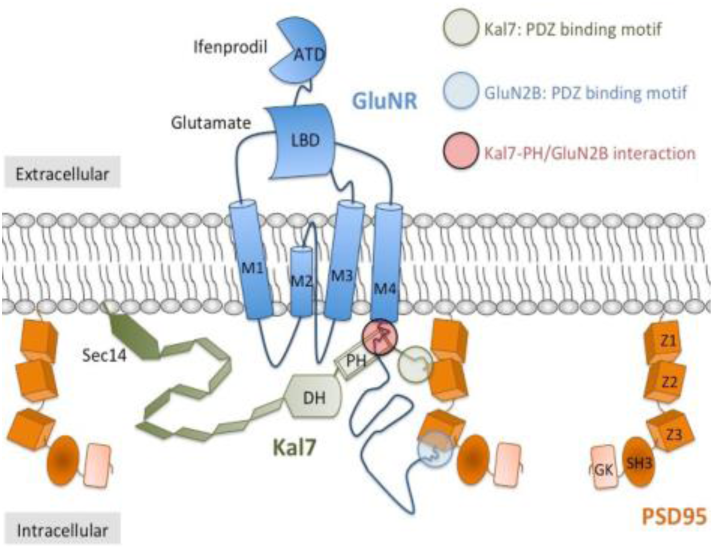

## Introduction

Genetic studies have confirmed a role for *KALRN* in schizophrenia, intellectual disability, attention deficit hyperactivity disorder, Alzheimer’s Disease (AD) and early onset coronary artery disease (Cahill et al., 2012; Deo et al., 2012; Hill, Hashimoto, & Lewis, 2006; Kushima et al., 2012; Lesch et al., 2008; Murray et al., 2012; Penzes & Jones, 2008; Russell et al., 2014; Wang et al., 2007; Wu et al., 2012; Youn, Jeoung, et al., 2007; Youn, Ji, Ji, Markesbery, & Ji, 2007). Kalirin-7 (Kal7), the major adult isoform, is concentrated in spines and co-fractionates with the postsynaptic density (PSD); Kal7 overexpression increases the linear density of spines and spine size (Ma et al., 2014; Ma, Wang, Ferraro, Mains, & Eipper, 2008; Penzes et al., 2001). Kal7 knockout mice exhibit a decrease in anxiety-like behavior, increased locomotor response to cocaine, increased self-administration of cocaine and a dramatic diminution of peripheral pain sensation (Kiraly, Lemtiri-Chlieh, Levine, Mains, & Eipper, 2011; Kiraly, Ma, Mazzone, Xin, & Mains, 2010; Kiraly et al., 2013; Lemtiri-Chlieh et al., 2011; Lu et al., 2015; Ma, Kiraly, et al., 2008; Mazzone, Larese, Kiraly, Eipper, & Mains, 2012). A mouse model of AD was used to demonstrate an essential role for Kal7 in the maintenance of normal spine density and cognitive behavior (Cisse et al., 2017).

Dendritic spines play a crucial role in learning and memory (Sheng & Kim, 2011). The ability of the Dbl homology (DH) domain of Kal7 to activate Rac1 has an essential role in controlling spine number, size and shape (Afroz, Parato, Shen, & Smith, 2016; Lu et al., 2015; Ma et al., 2014; Miller, Yan, Eipper, & Mains, 2013; Miller, Yan, Wu, et al., 2017). In addition to its guanine nucleotide exchange activity, the N-terminal Sec14 domain, spectrin repeat region and C-terminal PDZ binding motif of Kal7 play essential roles. In Kal7 knockout mice, hippocampal CA1 pyramidal neurons exhibit striking deficits in NMDA receptor-mediated currents, as indicated by a decrease in the NMDA/AMPA ratio (Ma, Kiraly, et al., 2008). Attenuation of NMDA receptor-dependent currents in Kal7 knockout mice is specific to GluN2B-containing receptors (Kiraly et al., 2011).

The plasticity phenomenon known as long-term potentiation (LTP) was discovered in 1973 (T. V. Bliss & Lomo, 1973; Larson & Munkacsy, 2015; Larson, Wong, & Lynch, 1986; Nugent, Hwong, Udaka, & Kauer, 2008) and is thought to underlie some forms of learning and memory. LTP is usually defined as an increase in synaptic strength lasting more than 1 hour. Two widely used patterns of stimulation to induce LTP in the hippocampus are high frequency stimulation (HFS: 100 Hz, often for 1-2 sec, and repeated a few times after 1-2 sec of quiescence) and theta burst stimulation (TBS; multiple trains of 100 Hz for 50 msec followed by 100-200 msec of quiescence) (T. V. P. Bliss & Collingridge, 1993; Larson & Munkacsy, 2015; Larson et al., 1986; Nugent et al., 2008). TBS is believed to disable feed-forward inhibition and better mimic *in vivo* hippocampal neuronal activity during learning paradigms and exploratory movements, and usually involves NMDA receptor activation and calcium fluxes (Larson & Munkacsy, 2015; Larson et al., 1986; Vanderwolf, 1969). Long-term depression (LTD) occurs in response to low frequency stimulation (LFS: e.g., 1 Hz, 15 min).

Since the specific sites through which Kal7 interacts with PSD95 and with the GluN2B subunit of the NMDA receptor have been mapped (Kiraly et al., 2011), we assessed the ability of synthetic Kal7 and GluN2B peptides to disrupt synaptic plasticity evoked by different stimulation paradigms. Our earlier field potential recordings of hippocampal CA1 pyramidal neurons demonstrated an essential role for Kal7 in LTP produced by TBS (but not by HFS) and in LTD produced by LFS. Although the stimulation paradigm needed to induce LTP in spinal projection neurons is distinctly different, inclusion of a peptide identical to the C-terminus of Kal7 in the recording patch pipette blocked LTP, mimicking the electrophysiological response observed in Kal7 knockout neurons (Lu et al., 2015).

## Materials and Methods

### Animal handling and slice preparation

All animal procedures were conducted using animal protocols approved by the University of Connecticut Institutional Animal Care and Use Committee. Postnatal day (P) 17-27 wildtype (WT) Swiss CD-1 mice (Charles River, Wilmington, MA) were anesthetized by 3.5% isoflurane inhalation, followed by decapitation. Kalirin-7 knockout mice (C57Bl/6) of the same age were treated identically (Ma, Kiraly, et al., 2008) (B6.129-*Kalrn*^*tm1Npl*^/J; Stock No:031465). Whole brains were removed and immersed in ice-cold slicing solution containing (in mM) 110 choline chloride, 2.5 KCl, 1.25 NaH_2_PO_4_, 25 NaHCO_3_, 0.5 CaCl_2_, 7 MgCl_2_, 25 D-glucose, 11.6 ascorbate, and 3.1 sodium pyruvate, equilibrated with 95% O_2_-5% CO_2_ (pH 7.3, 310 ± 5 mosmol/kg). Sagittal slices (300 μm) containing the hippocampus were cut with a Dosaka EM DTK-1000 vibratome (Kyoto, Japan) and transferred into an incubation chamber. Slices were then incubated for 15 minutes at 35**°**C in carboxygenated incubation solution containing (in mM) 125 NaCl, 2.5 KCl, 1.25 NaH_2_PO_4_, 25 NaHCO_3_, 0.5 CaCl_2_, 3.5 MgCl_2_, 25 D-glucose, 4 sodium lactate, sodium pyruvate, and 0.4 ascorbate (pH 7.3, 310 ± 5 mosmol/kg) before being transferred into the same incubation solution at room temperature. Slices were then individually transferred into a recording chamber (room temperature) fixed to the stage of an Olympus BX51WI upright microscope fitted with a X40 water-immersion lens (0.8 NA). The recording chamber was continuously perfused at 1.5 – 2 ml/min with carboxygenated artificial cerebrospinal fluid containing (in mM) 125 NaCl, 2.5 KCl, 1.25 NaH_2_PO_4_, 25 NaHCO_3_, 2 CaCl_2_, 2 MgCl_2_, and 25 D-glucose (pH 7.3, 305 ± 5 mosmol/kg).

### Electrophysiology

Whole cell recordings were obtained from pyramidal neurons in the CA1 region of the hippocampus. Pyramidal neurons were identified by their morphology and position under infrared differential interference contrast video microscopy. Patch electrodes (5-7 MΩ) were pulled from borosilicate glass capillaries using a Flaming/Brown P-97 micropipette puller (Sutter Instrument, Novato, CA). Pipette internal solution contained (in mM) 118.7 CH_3_O_3_SCs, 6.3 CsCl, 10 HEPES, 1 EGTA, 0.1 CaCl_2_, 1.5 MgCl_2_, 4 Na_2_-ATP, 0.3 Na-GTP, 10 di-tris-phosphocreatine and 5 QX-314 (pH 7.3, 290 ± 5 mOsm/kg). Cells were voltage clamped at −70 mV during recording. A bipolar tungsten electrode (1 MΩ, WPI) was positioned in the Schaffer collaterals emanating from the CA3 region of the hippocampus, approximately 100-150 μm lateral to the patched pyramidal neuron, in order to elicit electrically-evoked EPSCs (eEPSCs). Extracellular stimuli consisted of individual square-wave current pulses (170 μs, 4-30 μA) and were delivered every 20 seconds through a stimulus isolator (ISO-Flex; A.M.P.I.). Stimulation strength was set to a level that evoked 30-70% of the maximal response for each individual cell.

All electrical events were filtered at 2.9 kHz and digitized at > 6 kHz using a HEKA EPC9 amplifier and ITC-16 digitizer (HEKA Elektronik). Series resistance (*R*_s_) was compensated up to 20% at 100 μs lag. Input resistance (*R*_i_) was monitored with 10 mV (50 ms) hyperpolarizing steps at the end of each sweep. Cells were rejected from analysis if *R*_i_ fell below 50 MΩ or the holding current changed by > 15% during the course of an experiment.

### Peptides

All synthetic peptides were purchased from Biomatik (Wilmington DE); except where indicated, the amino-and carboxyl-terminal ends of the peptides were not modified. Peptide names and sequences are listed in **Table 1**. Peptides were reconstituted in the pipette internal solution and stored as concentrated stocks. Prior to recording experiments, each peptide was diluted in the pipette internal solution described above.

**Table 1.**
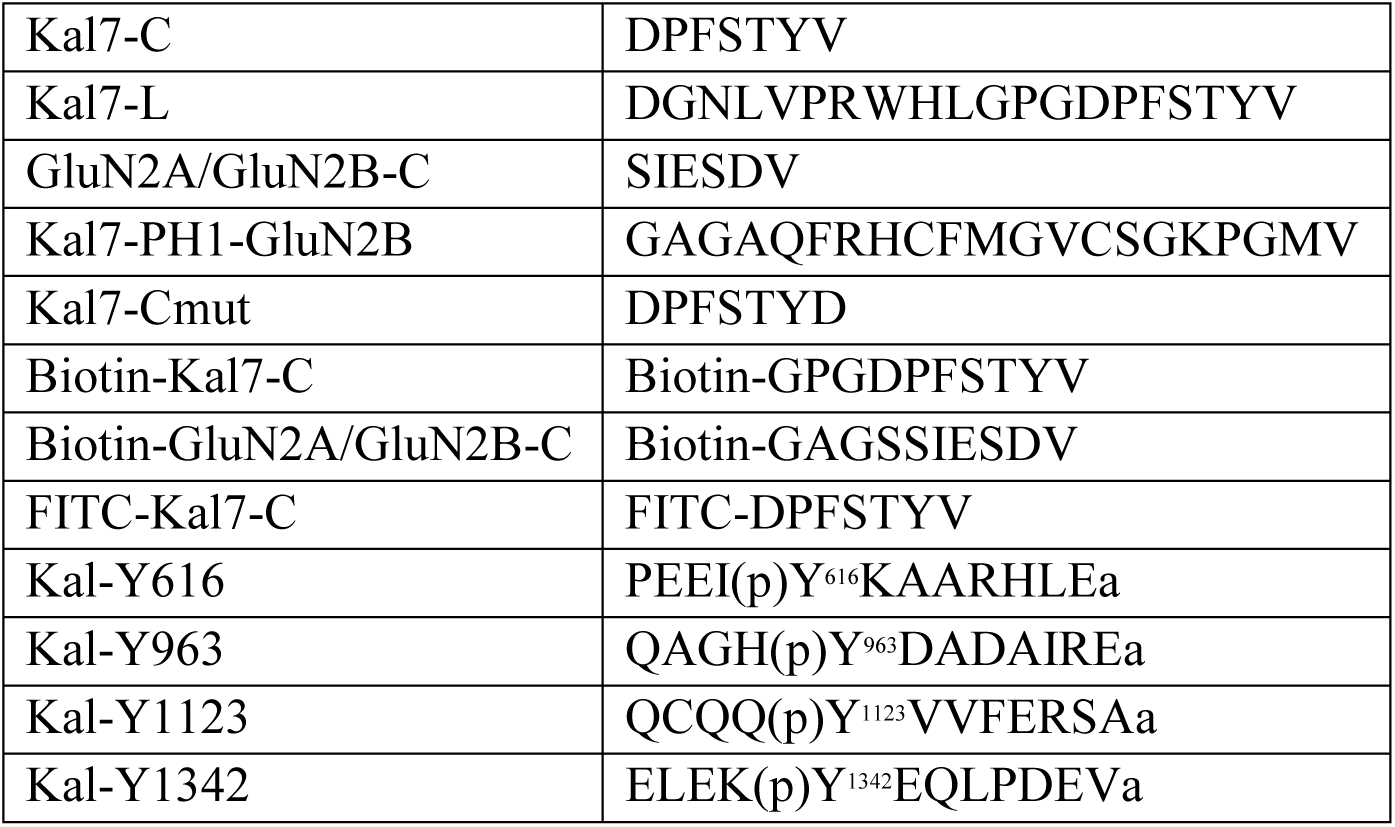
Peptides used for recording and binding studies

### Interactions of PSD-95 with Kal7C and GluN2B peptides

Biotinylated peptides were linked to StreptAvidin-and NeutrAvidin-Agarose (#29200 Thermo Scientific, Rockford IL) or SpeedBeed magnetic NeutrAvidin (GE Healthcare #78152104011150) per the manufacturer’s instructions. Vectors encoding PDZ1, PDZ2, PDZ3 and PDZ123 were kindly provided by Drs. Walkup and Kennedy (California Institute of Technology) and were expressed in BL21/DE3 bacteria (Walkup & Kennedy, 2014; Walkup et al., 2016). After induction of expression with isopropyl beta-D-1-thiogalactopyranoside, bacteria were harvested and extracted as described (Walkup & Kennedy, 2014; Walkup et al., 2016). Clarified lysates were incubated with Avidin-agarose bound biotinylated peptide (GluN2B-C or Kal7-C) in binding buffer (100Mm sodium phosphate, 50mM sodium chloride (pH 7.6), 0.1% TX100 buffer) and bound proteins were eluted in Laemmli SDS sample buffer; target peptides bound to ELISA plates were also tested as binding targets. Protein purity was verified by SDS-PAGE, followed by transfer to PVDF membranes, staining with Coomassie Brilliant Blue and Western blot analysis using a pan-MAGUK antibody (NeuroMab 75-028, clone 28/43; RRID AB_2292909) (Davis CA), a PSD-95 antibody (Upstate 05-427; RRID:AB_444362) (Lake Placid NY), and Kalirin antibodies JH2580 and JH2958 (Miller, Yan, Machida, et al., 2017) (RRID: 2801571; 2801572). Western blots were quantified in the linear range of exposures as described (Miller, Yan, Machida, et al., 2017). Similar experiments were conducted using mammalian PSD-95 protein (prepared by transfecting expression vectors pCMV-PSD95-flag [FLAG-tagged; Addgene, #15463] into HEK293 cells) bound to GluN2B-peptide-containing Agarose resins, as well as the binding of purified Kal7 [from baculovirus expression (Miller, Yan, Machida, et al., 2017)] paired with competing GluN2B peptides and control peptides (several biotinylated peptides from within the Kal7 sequence).

### Interactions of PSD-95 with Kal7 or Kal8

PDZ123 and control bovine serum albumin (BSA) were bound to UltraLink Biosupport resin (Thermo Scientific #53110) and to AffiGel 15 (Biorad # 1536051) per the manufacturers’ instructions. Resin was blocked with 1 mg/ml BSA for 15 min, washed in binding buffer, 0.6M Na-citrate (pH 7.4), binding buffer, then binding experiments were conducted and analyzed by Western blot (see below). For co-immunoprecipitation experiments, PSD95, Kal7, and Kal8 proteins were prepared by transfecting individual expression vectors into HEK293 cells: pCMV-PSD95-flag, an empty pFIV shuttle vector, pEAK-HisMyc-Kal7a, or pSCEP-Myc-Kal8 (Miller, Vishwanatha, Mains, & Eipper, 2015; Miller, Yan, Machida, et al., 2017). After 24 hours of transfection, HEK293 cells were rinsed in serum free medium (Rao, Zavala, Deb Roy, Mains, & Eipper, 2019), pelleted, and vortexed in ice-cold buffer containing 20mM TES, 10mM mannitol, 0.3mg/ml PMSF, 2μg/ml leupeptin, 2μg/ml pepstatin, 2 μg/ml benzamidine, 16μg/ml benzamidine, and 50μg/ml LBTI. Vortexed cell lysates were tumbled for 20 min at 4oC and centrifuged at 17,000g for 20 min; supernatant containing the expressed proteins of interest was collected. Protein samples of PSD95 and Kal7 or PSD and Kal8 were incubated at 4oC overnight with rabbit anti-Flag Antibody (#F7425 Sigma; RRID:AB_439687) in a final volume of 120μl 20mM TES, 10mM mannitol buffer containing 0.1% Triton X-100 (TMT buffer). After overnight incubation, samples were centrifuged at 17,000g for 10 min and supernatants were used for immunoprecipitation. Immobilized Protein A beads (10μl packed beads/ sample) (Repligen #10-2500-04) were rinsed 3 times with TMT buffer, and one time with 20mM TES, 10mM mannitol, 0.3mg/ml PMSF, 2μg/ml leupeptin, 2μg/ml pepstatin, 2 μg/ml benzamidine, 16μg/ml benzamidine, and 50μg/ml LBTI before addition of samples. After samples and protein A beads were tumbled at 4oC for 1 hour, unbound proteins were removed and beads were washed twice with 0.5ml TMT buffer and once with 0.5ml TM buffer (20mM TES, 10mM mannitol). Bound proteins were eluted by boiling into 2x Laemmli sample buffer. For each sample, incubation of equal amounts of protein with a nonimmune rabbit IgG antibody was used as a negative control for nonspecific binding. An aliquot of each sample was saved as input, and equal amounts of input were used. Samples were analyzed on 4% to 15% gradient gels (Criterion TGX; BioRad) after transfer to a polyvinylidene difluoride membrane. Antibodies used included PSD-95 (#75028 Neuromab), FLAG monoclonals [Proteintech 66008-3-Ig, RRID:AB_2749837; Sigma M2 #F1804, RRID:AB_262044], rabbit anti-Flag Antibody (Sigma #F7425, RRID:AB_439687) and affinity-purified CT302 (Yan, Eipper, & Mains, 2015) (RRID:AB_2801573) for the detection of Kal7 and Kal8). Immunoblots were visualized using SuperSignal West Pico PLUS Chemiluminescent substrate (Thermo Scientific) and quantified in the linear range of signals with the SynGene Pxi4 imaging system.

### Experimental design and statistical analysis

Group data are reported as mean ± SE. Statistical comparisons were made using one-way ANOVA and Dunnett’s multiple comparison test or Student’s paired t-test for post-hoc comparison. p < 0.05 was taken as a statistically significant effect. Long term potentiation and depression are always compared to untreated cells (no peptide) subjected to the same electrical stimulation pattern.

## Results

In our previous electrophysiological analyses of CA1 pyramidal neurons, we used field potential recordings in acute slices to demonstrate an important role for Kal7 in the LTP that occurs in response to theta burst stimulation (TBS) and in the LTD that occurs in response to prolonged 1 Hz stimulation (low frequency stimulation, LFS). Strikingly, the absence of Kal7 did not diminish the NMDA receptor-independent LTP observed in response to 200 Hz/2sec tetanic stimulation in the presence of CPP (Kiraly et al., 2011; Lemtiri-Chlieh et al., 2011; Ma, Kiraly, et al., 2008). Pharmacological and biochemical studies lent support to the conclusion that Kal7, which interacts with both PSD95 and GluN2B, plays a unique role in GluN2B-dependent signaling (Kiraly et al., 2011; Lemtiri-Chlieh et al., 2011; Ma, Kiraly, et al., 2008; Penzes et al., 2001).

### Synaptic plasticity deficits observed in CA1 pyramidal neurons lacking Kal7 using patch clamp electrophysiology

With the goal of acutely disrupting the interactions of Kal7 with PSD95 or with GluN2B by injecting synthetic peptides into wild type neurons, we first verified that whole cell patch clamp electrophysiology revealed similar deficits in LTP and LTD in CA1 pyramidal neurons in slices prepared from Kal7 knockout mice (**Fig. 1**). Two different induction protocols were used, both under voltage clamp conditions: (1) 7 train TBS and (2) HFS (two 100 Hz bursts for 2 seconds, separated by 20 seconds). For the 7 train TBS, each TBS train contained 10 bursts (200 ms interburst interval), each burst consisted of 5 stimuli at 100 Hz. Each train was delivered with a 5 s intertrain interval. As observed using field potential recordings (Lemtiri-Chlieh et al., 2011), whole cell patch clamp recordings revealed the inability of Kal7 knockout CA1 pyramidal neurons to exhibit LTP in response to administration of 7 train TBS (**Figs. 1A** and **B**). In contrast, patched Kal7 knockout neurons subjected to HFS exhibited robust LTP (**Figs. 1A** and **C**), as did WT neurons stimulated with both TBS (p = 0.02) and HFS (p = 0.04) stimulation paradigms (not shown). Whole cell patch recordings from Kal7 knockout neurons exposed to LFS (1 Hz) revealed their failure to exhibit LTD (**Fig. 1D**), as observed using field recordings (Kiraly et al., 2011; Lemtiri-Chlieh et al., 2011). Thus the use of whole cell patch clamp to deliver synthetic peptides designed to disrupt acutely the interactions of Kal7 with PSD95 or GluN2B and assess the effects of these peptides on the efficacy of specific plasticity induction paradigms is justified.

**Figure 1.**
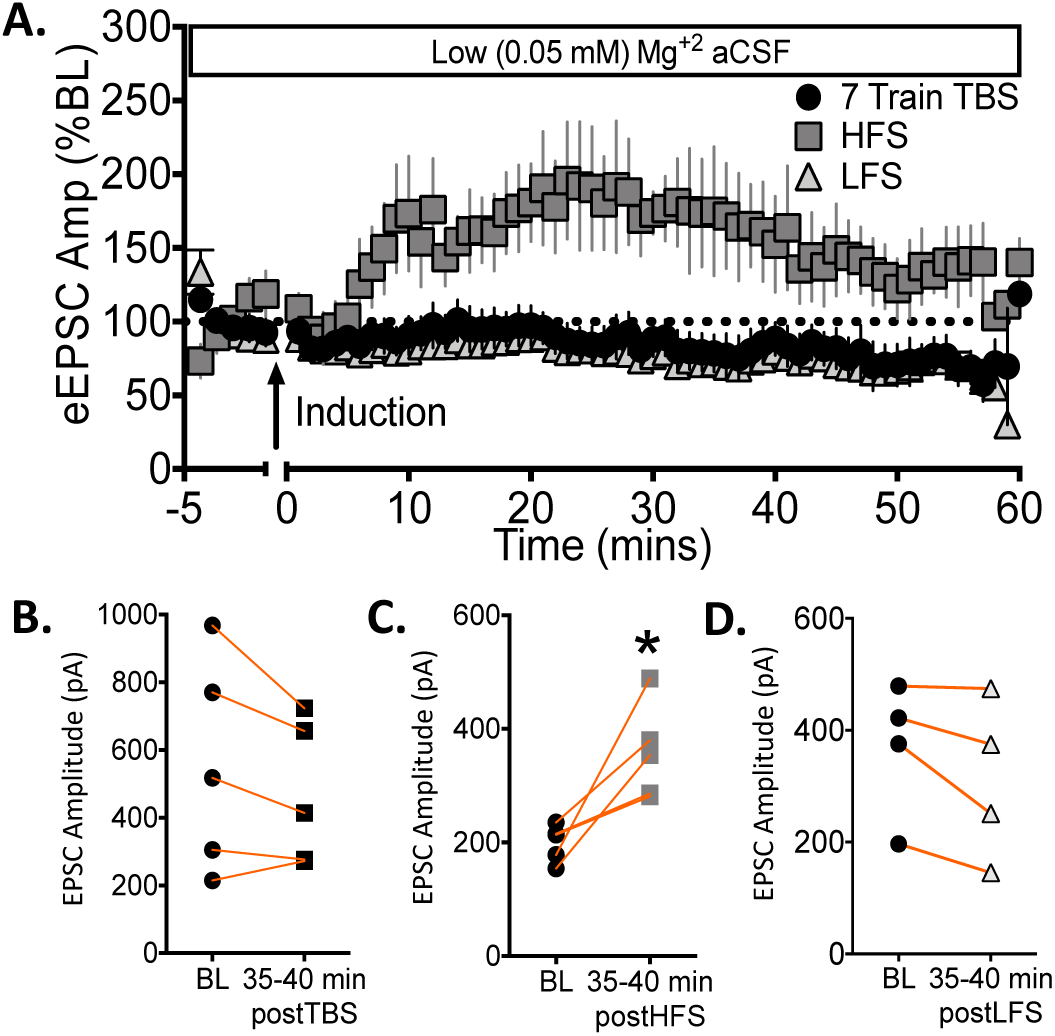
TBS, but not HFS, LTP is disrupted in CA1 hippocampal pyramidal neurons of Kal7 knockout mice. (A) Group time course of eEPSC amplitude of CA1 pyramidal neurons in Kal7 knockout mice. Two induction protocols are shown. Cells subjected to 7 train TBS (black circles) do not exhibit LTP. However, cells subjected to 100 Hz HFS showed robust LTP (p=0.02). (B) Before and after plot of pyramidal neurons in CA1 of hippocampus in Kal7 knockout mice show no changes in eEPSC amplitude (pA) following induction with 7 train TBS between baseline and 35 – 40 mins post-TBS. (C) Before and after plot of pyramidal neurons in CA1 of hippocampus in Kal7 knockout mice shows an increase in amplitude (pA) after 100 Hz HFS (p=0.02). (D) Before and after plot of pyramidal neurons in CA1 of hippocampus in Kal7 knockout mice shows no change in eEPSC amplitude (pA) or area (pA * ms) after 1 Hz LFS. For 7 train TBS induction protocol, 5 cells recorded from 2 mice (1 male; 1 female). For HFS induction protocol, 5 cells recorded from 2 mice (1 male; 1 female). For 1 Hz LFS induction protocol, 4 cells recorded from 2 mice (1 male; 1 female).

### Acute disruption of specific Kal7/PSD95/GluN2B interactions with synthetic peptides blocks distinct forms of synaptic potentiation

Studies from many laboratories have provided detailed information about the roles of GluN2B, PSD95 and Kal7 in synaptic function (T. V. P. Bliss & Collingridge, 1993; Larson & Munkacsy, 2015; Larson et al., 1986; Nugent et al., 2008). All three proteins are enriched at the PSD (Grant, 2006; Kiraly et al., 2011; Sheng & Hoogenraad, 2007; Sheng & Kim, 2011). Quantitative studies indicate that the average PSD contains about 300-400 PDZ domain family members, with PSD95 by far the most prevalent. Proteins with PDZ-binding motifs known to recognize PSD95 are enriched at the PSD, with about 20 copies of Kal7, 60 NMDA receptor subunits, 60 AMPA receptor subunits and 20 mGluR’s, all of which have PDZ-binding motifs (Grant, 2006; Kiraly et al., 2011; Sheng & Hoogenraad, 2007; Sheng & Kim, 2011). In addition, the PH1 domain of Kal7 interacts with the final juxtamembrane domain of GluN2B (Kiraly et al., 2011) (**Fig. 2A**).

**Figure 2.**
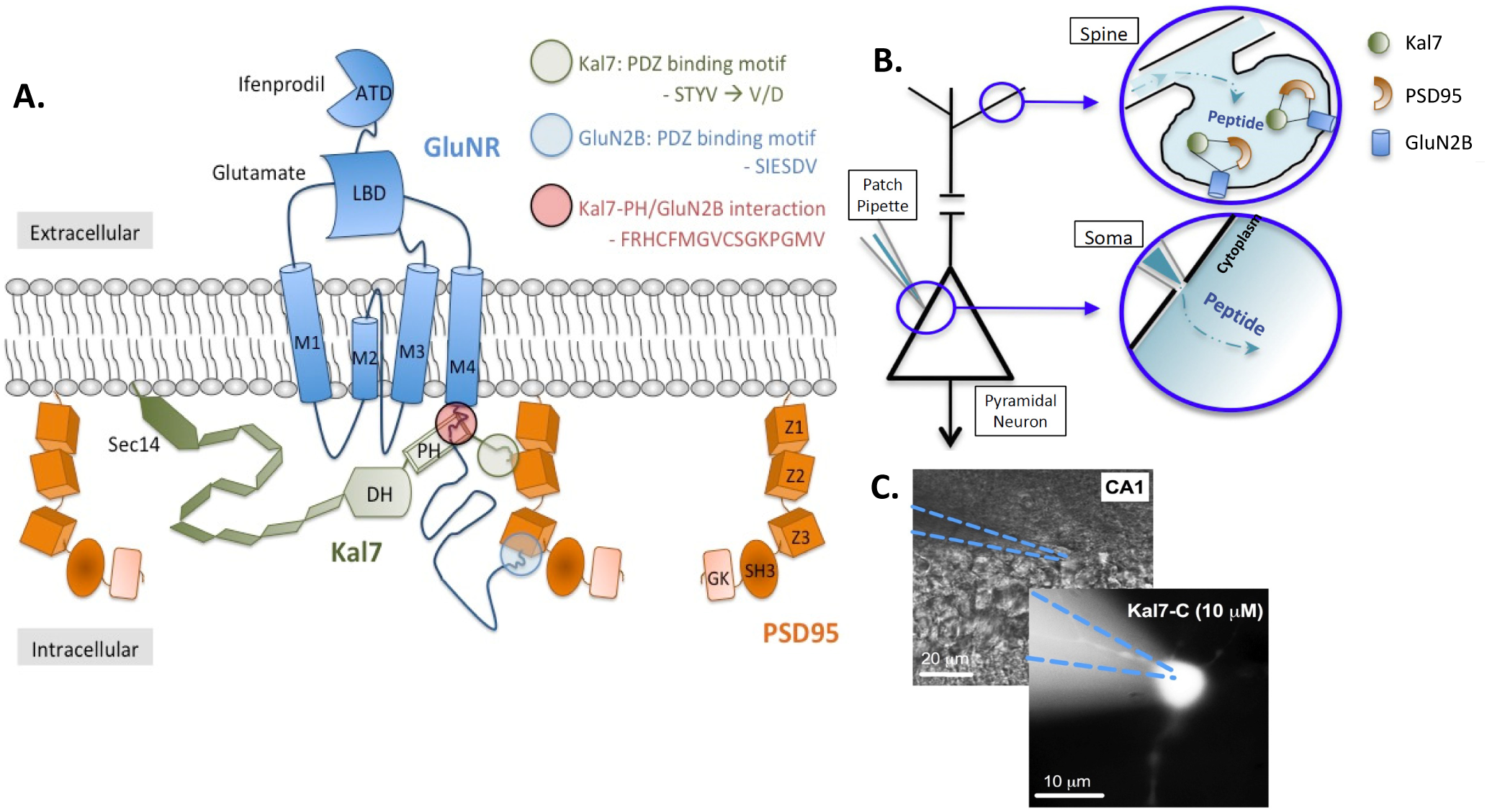
Arrangement of postsynaptic proteins and delivery of synthetic peptide via patch pipette. (A) Cartoon rendering of a GluN2B-subunit containing NMDA receptor (blue) showing putative sites of interaction with Kal7 (green) and PSD95 (orange). Color-coded circles indicate target binding sites of synthetic peptides for (1) Kal7-C, (2) GluN2B and (3) GluN2B-Kal7 within the postsynaptic terminal. (B) Schematic illustrating the intracellular delivery of synthetic peptides under whole cell patch clamp conditions. Synthetic peptides are diluted to desired working concentrations in the intracellular recording solution and loaded into the recording pipette. Cells are allowed to fill for 10 minutes, during which time the intracellular recording solution containing the diluted peptide is distributed within the soma and dendrites and presumably, the dendritic spines (magnified fields of view). (C) Kal7-C tagged with FITC was visualized in a neuron situated in CA1 of mouse hippocampus after 10 minutes of whole cell recording mode. The patched neuron can be seen under DIC and under epifluorescence, which shows the soma and processes filled with FITC-tagged Kal7-C peptide.

Our goal was to determine whether acute intracellular administration of peptides designed to mimic the C-terminal PDZ-binding motifs of Kal7 (Kal7-C) and GluN2B (GluN2B-C) and the PH-domain of Kal7, which interacts with the juxtamembrane domain of the GluN2B subunit (Kal7-GluN2B), had any effect on three different types of synaptic plasticity. Peptides diluted into intracellular recording solution were injected into CA1 hippocampal pyramidal neurons before whole cell patch clamp electrophysiological recordings were begun. Using a FITC-tagged Kal7-PDZ peptide, we found that significant filling of the patched CA1 hippocampal neuron occurred within 10 min of breaking into the cell (**Figs. 2B**,**C**).

Control cells where no peptide was added to the intracellular recording solution showed robust LTP following induction with 7 train TBS (**Fig**.**3A**). We first asked whether acute disruption of the Kal7-PSD95 interaction blocked LTP induced by 7 train TBS stimulation (**Fig**.**3A**). When used at a concentration of 5 or 10 μM, Kal7-C eliminated TBS-induced LTP. Injection of 1 μM Kal7-C was without effect on TBS-induced LTP.

### Control peptides do not interfere with LTP

As noted above, a lot of proteins in cells have PDZ binding motifs. To test for specificity and toxicity, the C-terminal Valine residue of Kal7-C was replaced by Aspartic Acid (STYV **→** STYD) (Kal7-C_mut_), destroying the PDZ-binding motif (Songyang et al., 1997). Following the injection of Kal7-C_mut_, 7 train TBS elicited LTP that resembled the LTP observed in control cells at 35 – 40 minutes post-induction at a concentration of either 1 μM (p = 0.01) or 10 μM (**Fig**.**3B**). While the effect of this single amino acid change supports the specificity of the peptide effect, the PDZ-binding motif of Kal7 may interact with multiple proteins in addition to PSD95.

In an effort to elucidate potential effects of the Kal7-C short peptide on baseline glutamatergic transmission (spontaneous and electrically-evoked), we collected data from a separate cohort of cells that included single evoked excitatory postsynaptic currents (eEPSCs) and spontaneous excitatory postsynaptic currents (sEPSCs) over the course of the first 10 minutes after entering the whole cell recording configuration. Over a 20 minute period, eEPSC **(Fig. 3D**) and sEPSC (**Fig. 3E**) amplitude and area remained stable from the time of break-in onward. This result is consistent with observations made in hippocampal slices from WT and Kal7 knockout mice using field recordings (Lemtiri-Chlieh et al., 2011).

**Figure 3.**
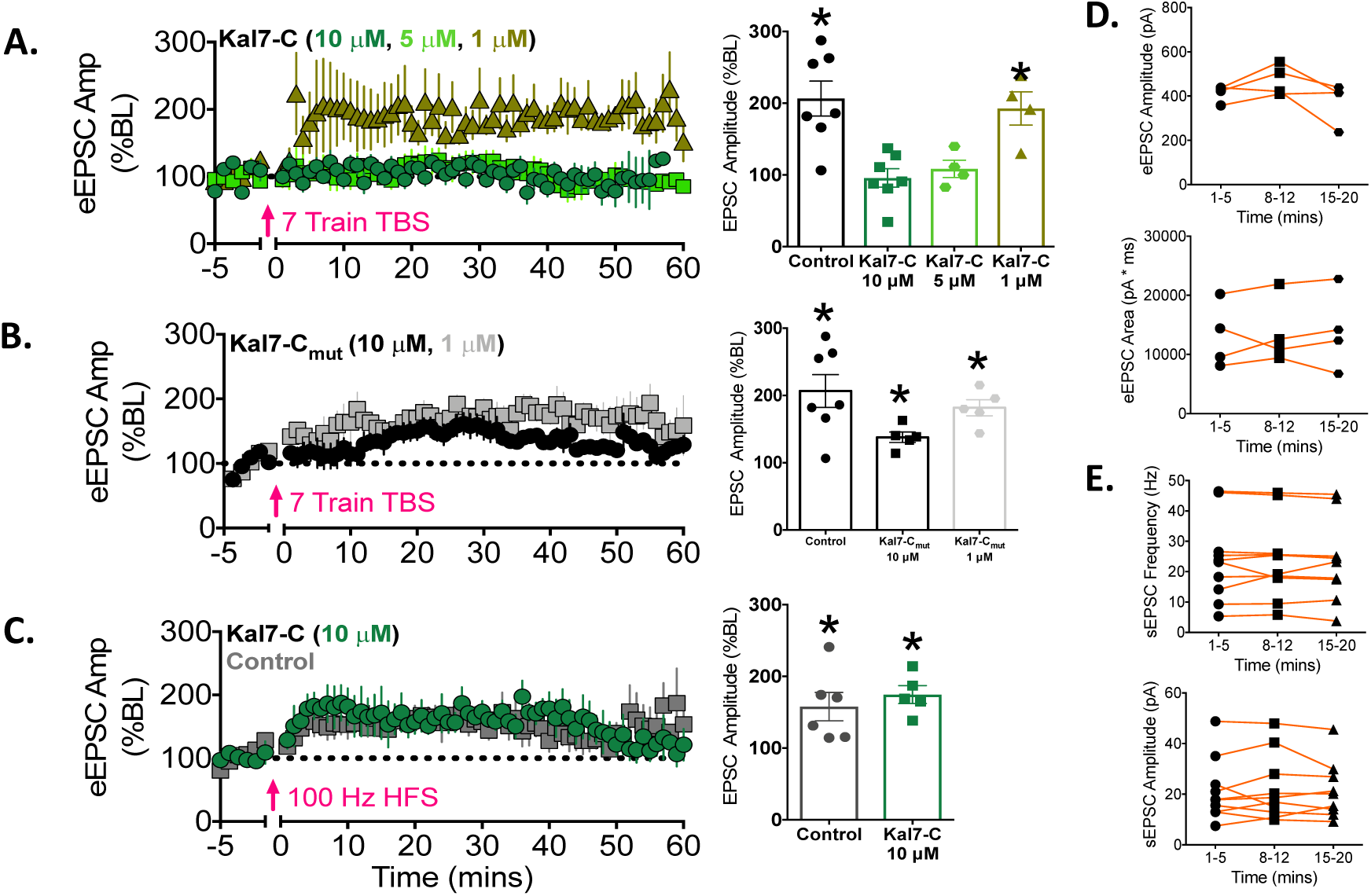
Acute disruption of Kal7/PSD95 interaction blocks LTP induced by TBS, but not by HFS, in CA1 hippocampal pyramidal neurons. (A) Left; Group time course plot of pyramidal neurons in CA1 of hippocampus filled with Kal7-C at 10 μM (dark green), 5 μM (light green) and 1 μM (olive) showing baseline and post-induction using 7 train TBS. Control LTP was 155% (p=0.02 vs. baseline); Kal7-C peptide at 10 μM and 5 μM blocked induction of LTP, while blockade was not observed at 1 μM Kal7-C peptide (p=0.05). Right; Grouped and individual data for each treatment at 35 – 40 mins post-induction. Asterisks denote statistical significance of within-group comparisons. (B) Left; Group time-course of pyramidal neurons employing intracellular Kal7-C_mut_ at 10 μM (black circles) and 1 μM (light grey squares) shows no disruption of LTP following 7 train TBS induction protocol. Right; Group means and individual data. Asterisks denote statistical significance for within-group comparisons. (C) Left; Group time-course showing that control cells (grey squares) and cells filled with Kal7-C at 10 μM (green circles) express LTP following 100 Hz HFS induction. Right; Group means. For control cells subjected to 7 train TBS, 7 cells recorded from 3 mice (2 male; 1 female). (D) Time course of neurons filled with Kal7-C peptide (10 µM) showing no change in eEPSC amplitude or area over 20 minutes (4 cells; 2 mice (1 male, 1 female)). (E) Time course of neurons filled with Kal7-C peptide (10 µM) showing no change in sEPSC frequency or amplitude over 20 minutes (10 cells; 3 mice (2 male, 1 female)). Control cells subjected to 7 train TBS in panel A are the same cells represented in panel B. In addition, control cells subjected to 7 train TBS are also presented in Figures 4 and 5 below. Control cells subjected to 100 Hz HFS, 6 cells recorded from 3 mice (2 male; 1 female). Control cells subjected to 100 Hz HFS are also presented in Figures 4 and 5. For cells filled with Kal7-C 10 μM, 7 cells recorded from 4 mice (2 male, 2 female); Kal7-C 5 μM, 4 cells recorded from 3 mice (1 male, 2 female); Kal7-C 1 μM, 4 cells recorded from mice (2 male, 1 female). For cells filled with Kal7-C_mut_ 10 μM, 5 cells recorded from 2 mice (1 male, 1 female); Kal7-C_mut_ 1 μM, 5 cells recorded from 2 mice (1 male, 1 female).

**Figure 4.**
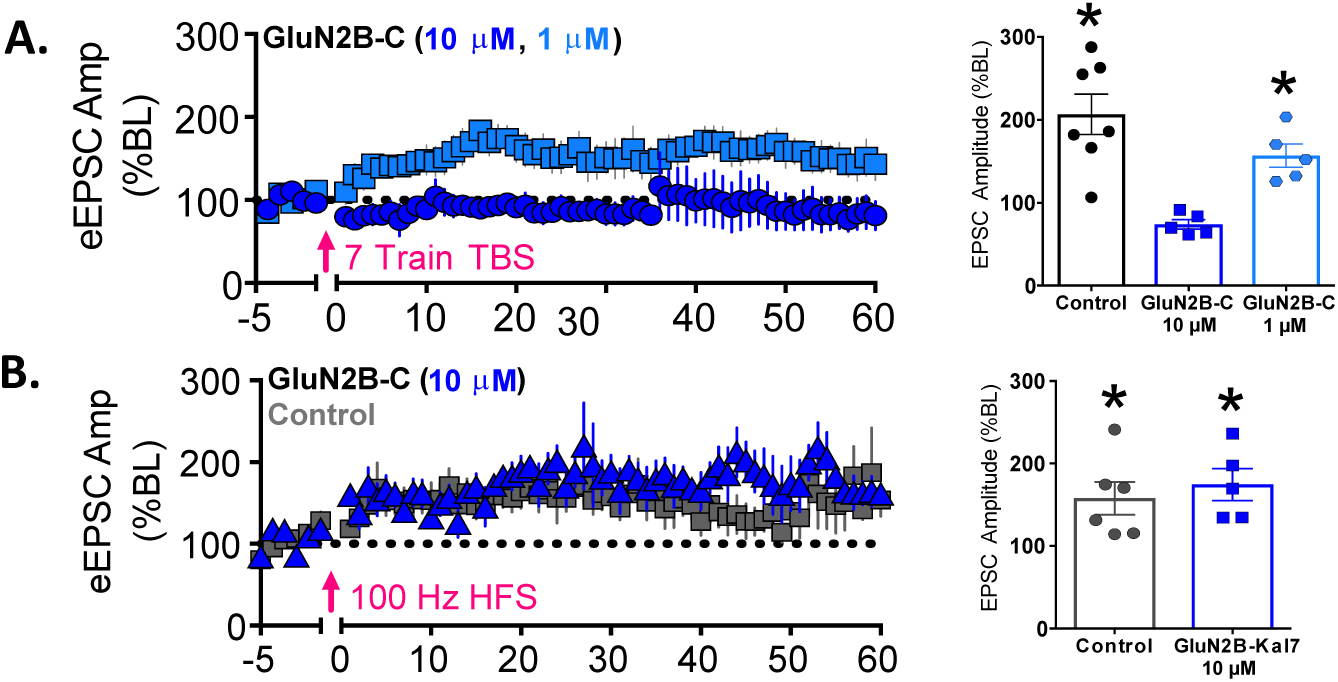
Acute disruption of GluN2B/PSD95 interaction blocks LTP induced by TBS, but not by HFS, in CA1 hippocampal pyramidal neurons. (A-B) Group time-course plots show pyramidal neurons subjected to (A) 7 train TBS after filling with 10 μM or 1 μM GluN2B-C peptide and subjected to (B) 100 Hz HFS after filling with 10 μM GluN2B-C peptide. Column graphs show group means at 35 – 40 mins post-induction. Asterisks denote statistical significance for within-group comparisons. For cells filled with GluN2B-C 10 μM subjected to 7 train TBS, 5 cells recorded from 2 mice (1 male, 1 female). For cells filled with GluN2B-C 1 μM subjected to 7 train TBS, 5 cells recorded from 3 mice (1 male, 2 female). For cells filled with GluN2B-C 10 μM subjected to 100 Hz HFS, 5 cells recorded from 2 mice (1 male, 1 female).

**Figure 5.**
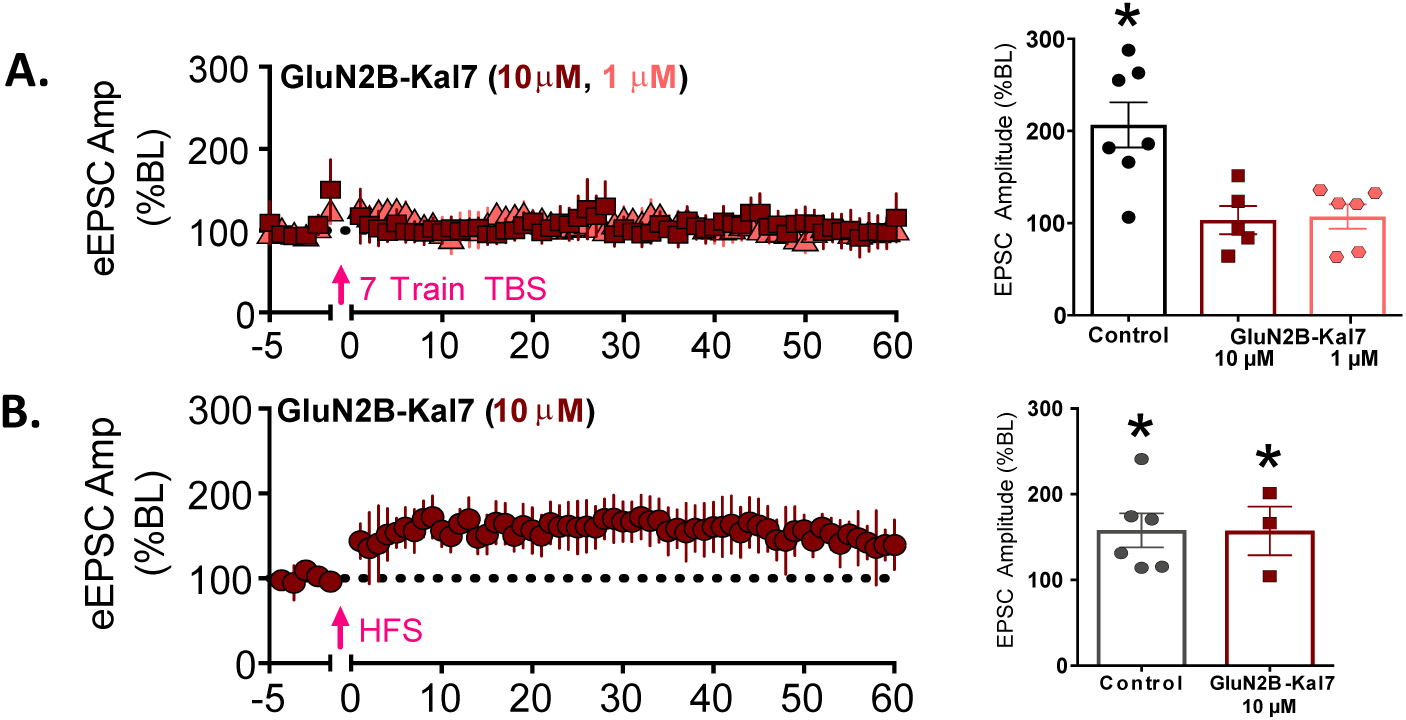
Acute disruption of GluN2B/Kal7 interaction blocks LTP induced by TBS, but not by HFS, in CA1 hippocampal pyramidal neurons. (A-B) Group time-course plots show pyramidal neurons subjected to (A) 7 train TBS after filling with (A) 10 μM or 1 μM GluN2B-Kal7 peptide and subjected to (B) 100 Hz HFS after filling with 10 μM GluN2B-Kal7 peptide. Histograms show group means at 35 – 40 mins post-induction. Asterisks denote statistical significance for within-group comparisons. For cells filled with GluN2B-Kal7 10 μM subjected to 7 train TBS, 5 cells recorded from 2 mice (1 male, 1 female). For cells filled with GluN2B-Kal7 1 μM subjected to 7 train TBS, 6 cells recorded from 3 mice (2 male, 1 female). For cells filled with GluN2B-Kal7 subjected to 100 Hz HFS, 3 cells recorded from 2 mice (1 male, 1 female).

**Figure 6.**
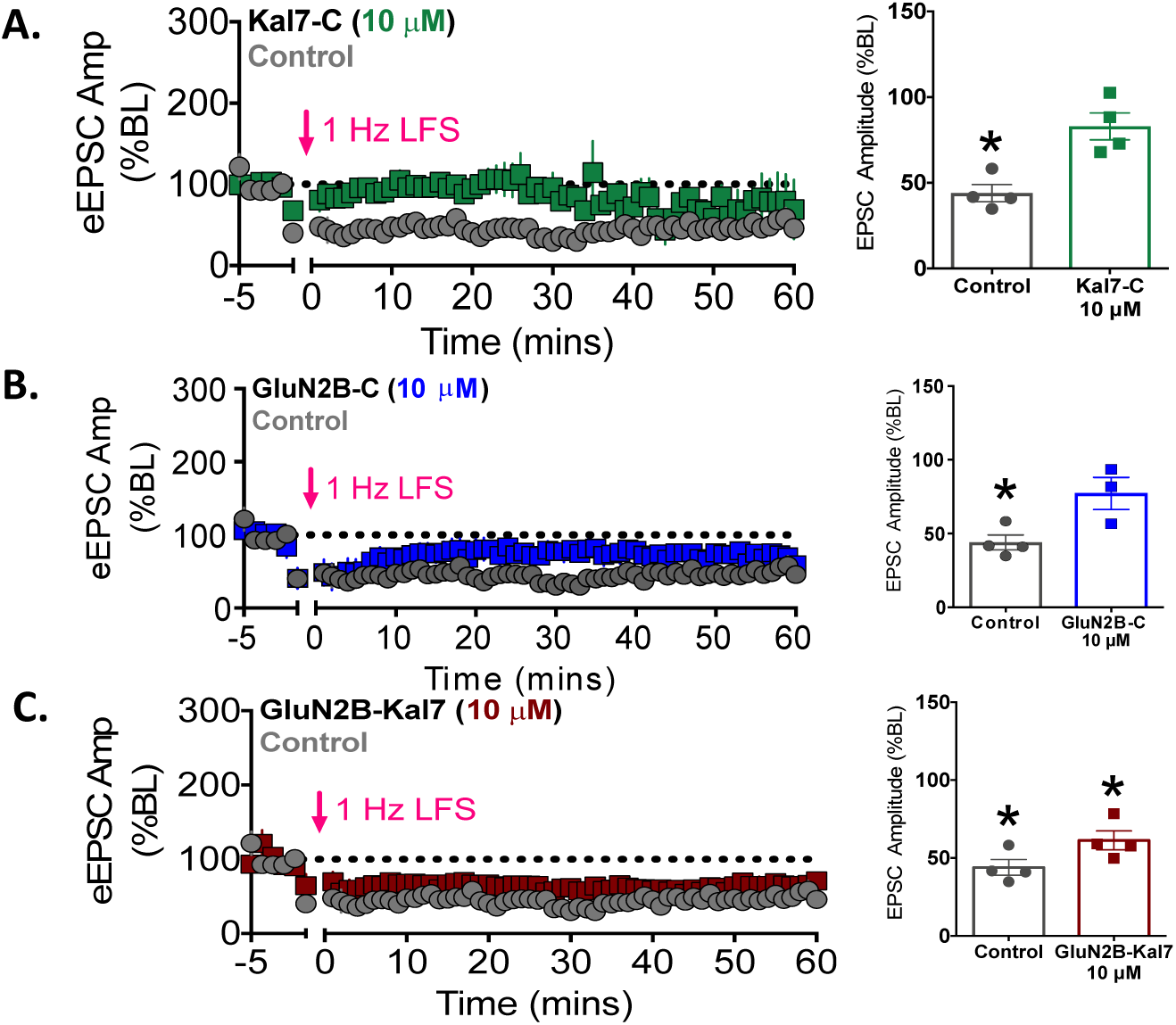
Effects of acute disruption of Kal7/PSD95, GluN2B/PSD95 or GluN2B/Kal7 on LTD in CA1 hippocampal pyramidal neurons. (A-C) Group time-course plots showing cells filled with (A) Kal7-C peptide, (B) GluN2B-C peptide or (C) GluN2B-Kal7 peptide at 10 μM before being subjected to 1 Hz LFS. The Kal7/PSD95 peptide blocked LTD in pyramidal neurons situated in CA1 of the hippocampus (A). The GluN2B/PSD95 peptide did not affect the early phase of LTD (0 to 5 min), but appears to block LTD at longer post-induction times (>20 min) (B). The GluN2B-Kal7 peptide had no significant effect on LTD at any time following 1 Hz LFS (C). For control cells without peptide present in the internal recording solution and subjected to 1Hz LFS, 4 cells recorded from 3 mice (2 male, 1 female). Control cells without peptide apply to all panels. For cells filled with Kal7-C 10 μM, 4 cells recorded from 2 mice (1 male, 1 female). For cells filled with GluN2B-C 10 μM, 3 cells recorded from 2 mice (1 male, 1 female). For cells filled with NR2B-Kal7 10 μM, 4 cells recorded from 2 mice (1 male, 1 female).

**Figure 7.**
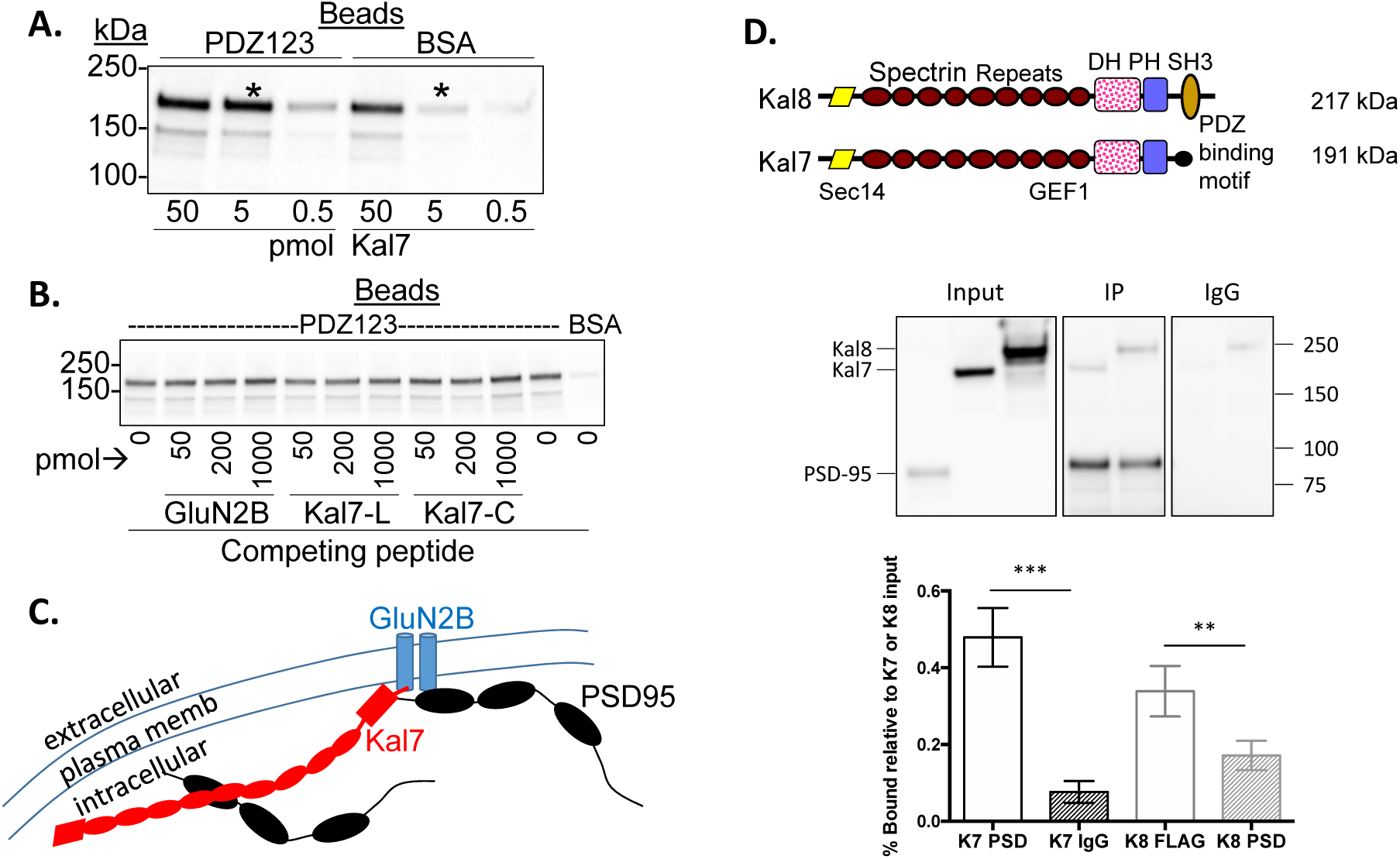
PSD-95 associates with both Kal7 and Kal8 *in vitro*. (A) UltraLink Biosupport resin containing 1 nmol (1000 pmol) PDZ123 or BSA was incubated with the indicated amounts of purified Kal7 for one hour at 4oC, washed in binding buffer and analyzed by Western. (B) PDZ123 beads as in (A) were incubated with 5 pmol purified Kal7 plus the indicated amounts of potential interfering peptides, followed by Western analyses of the beads. (C) Simplified model of the multiple interactions possible at the postsynaptic terminal among Kal7, PSD95 and GluN2B. (D) Immunoprecipitation (IP) was performed on mixtures of FLAG-tagged PSD-95, Kal7 (K7), and Kal8 (K8) protein extracts from transfected HEK293 cells using FLAG rabbit antibody or an equal amount of nonimmune rabbit IgG. Co-IP revealed interactions between PSD-95 with both Kal7 and Kal8, regardless of the presence of a COOH-terminal PDZ-binding domain; IgG controls show no significant binding of protein. Co-IP FLAG antibody of Kal7 or Kal8 with PSD-95 is significantly greater than control IgG antibody; students paired t-test, p=0.0008 [***] and p= 0.0031 [**], respectively. Unpaired t-test comparing bound Kal7 FLAG with bound K8 FLAG was not statistically significant, p= 0.189. N=7.

Although mice lacking Kal7 throughout gestation and maturation fail to exhibit LTP in response to TBS, HFS stimulation produces LTP that is equivalent to that observed in wildtype mice (Lemtiri-Chlieh et al., 2011). We next asked whether the acute presence of Kal7-C peptide affected LTP in response to HFS stimulation (**Fig**.**3C**). Injection of 10 μM Kal7-C peptide had no effect on the ability of CA1 hippocampal neurons to exhibit LTP in response to HFS (**Fig**.**3C**). This result is consistent with observations made using field recordings and Kal7 knockout mice (Lemtiri-Chlieh et al., 2011).

### Acute disruption of GluN2A/B←→PSD95 interaction blocks TBS, but not HFS, induced LTP

Biochemical and pharmacological analyses of Kal7 knockout mice have provided strong support for the importance of GluN2B in the actions of Kal7. Both GluN2B and Kal7 terminate with PDZ binding motifs known to interact with PSD95; whether the two motifs bind simultaneously to PSD95 or interfere with the other’s binding to PSD95 is unknown. In addition to their shared interaction with PSD95, the PH1 region of Kal7 interacts directly with GluN2B (**Fig**.**2A**).

As expected, administration of 7 train TBS stimulation after bath application of CPP (5 μM), a selective NMDA receptor antagonist, blocked LTP in all cells examined (not shown). We first explored the role of the GluN2A/GluN2B subunit in TBS induced LTP by injecting a 6-residue peptide identical to the C-terminus of GluN2A/GluN2B (**Fig**.**4**). Using GluN2A/GluN2B-C at 10 μM, TBS was unable to elicit LTP (**Fig**.**4A**). When the concentration of GluN2A/GluN2B in the intracellular recording solution was decreased to 1 μM, 7 train TBS elicited LTP that was indistinguishable from control cells at 35 – 40 minutes post-induction (**Fig**.**4A**). Dilution of the Kal7-C peptide (**Fig**.**3A**) suggests that the binding motifs of GluN2A/GluN2B and Kal7 exhibit similar affinities for their target PDZ domains.

We next tested the effect of the GluN2A/GluN2B peptide on LTP induced by HFS. Following the injection of 10 μM GluN2A/GluN2B-C, HFS resulted in LTP that was comparable to that observed in control cells (**Fig**.**4B**). As seen with the Kal7-C peptide, HFS stimulated LTP was not affected by the GluN2A/GluN2B-C peptide.

### Acute disruption of GluN2B-Kal7 interaction blocks TBS, but not HFS, induced LTP

Our data suggest that both the Kal7-C and GluN2A/GluN2B peptides interact with PSD95 or another PDZ-domain protein in the dendritic spine within minutes of breaking into the cell, in order to block LTP following the 7 train TBS induction protocol. For this reason, we decided to probe the interaction between Kal7 and the juxta-membrane domain of GluN2B, at the GluN2B-specific interaction site described by Kiraly and colleagues (Kiraly et al., 2011) (**Fig**.**2A**). At a concentration of 10 μM, the presence of Kal7-GluN2B peptide in the intracellular recording solution blocked 7 train TBS-dependent LTP in all cells examined (p = 0.43) (**Fig**.**5A**). Even when tested at 1 μM, the Kal7-GluN2B peptide still blocked 7 train TBS-dependent LTP, suggesting that the Kal7/GluN2B interaction is more readily disrupted by exogenous peptide than the PDZ-domain interactions mediated by Kal7-C and GluN2A/GluN2B-C (p = 0.99).

As for the PDZ-binding motif peptides, HFS-dependent LTP was not inhibited by the presence of Kal7-GluN2B at 10 μM, further reinforcing the evidence that HFS induction may elicit LTP independently of Kal7 and GluN2B-subunit containing NMDA receptors (p = 0.05) (**Fig**.**5B**).

### Acute disruption of Kal7←→GluN2A/B←→PSD95 interactions also alters LTD

Mice lacking Kal7 throughout gestation and maturation do not exhibit LTD in response to LFS (Lemtiri-Chlieh et al., 2011). We next tested whether the acute addition of Kal7-C peptide blocked LTD in response to LFS stimulation (**Fig**.**6**). LFS produced the expected long-lasting depression of eEPSC amplitude (to approximately 50%) in control neurons (**Fig**.**6A**). The injection of Kal7-C peptide in the recording pipette abolished LTD for at least 30 min; there was noticeable recovery of LTD toward the end of the hour-long recording session. However, even at 35-40 minutes post-induction, eEPSC amplitude was not statistically different from baseline. Thus, injection of Kal7-C peptide disrupted LFS-induced LTD seen in control neurons, which corroborates data from Kal7 KO mice.

When LFS induced LTD was tested in the presence of GluN2A/B peptide in the recording pipette (**Fig**.**6B**), LTD was initially unaffected; with increasing time, the presence of GluN2A/B gradually blocked the maintenance of LTD (beyond 20 min).

Lastly, we examined the effect of the Kal7-GluN2B peptide (which was remarkably potent at blocking LTP [**Fig**.**5A**]) on LFS-induced LTD (**Fig**.**6C**). Unlike the Kal7-C and GluN2B peptides, cells filled with Kal7-GluN2B (10 μM) showed no differences in LTD compared to control cells at any time post-induction and expressed robust LTD following LFS. The differences in the test peptide blocking results for LTP and LTD suggest important differences in the fundamental underlying causes of these long term changes in plasticity (Zhou et al., 2018). Specifically, GluA1 (which ends …ATGL; a Class 1 PDZ motif, as are the peptides being tested here) is crucial for LTP, while GluA2 (which ends …SVKI, a Class 2 PDZ motif) is crucial for LTD (Zhou et al., 2018).

### Biochemical examination of interactions of the test peptides and proteins with PSD95

PDZ domains are 80-90 residues in length; there are 300-400 such domains in over 150 proteins (Caillet-Saguy et al., 2015; Houslay, 2009). PDZ binding motifs typically consist of only 3-4 primary residues, primarily at the C-terminus of proteins (Caillet-Saguy et al., 2015). The initial interaction of a PDZ binding motif with a PDZ domain is generally followed by an induced fit (Chi et al., 2009). PDZ-binding motifs are common; over a third of the 70+ Rho-GEFs contain PDZ-binding motifs, with similar numbers in *Drosophila* and *C*.*elegans*, but no known PDZ domains in yeast (Garcia-Mata & Burridge, 2006). A number of viruses capitalize upon their PDZ-binding motifs to spread within infected cells (Caillet-Saguy et al., 2015; Pim et al., 2012).

In order to address the biochemical interaction of Kal7 and GluN2B, more specifically if GluN2B and Kal7 compete for binding sites on PSD-95, the initial biochemical attacks utilized the same interfering peptides used for the electrophysiological experiments. Because Kal7 and GluN2B both have PDZ binding motifs that interact with PSD-95, the first approach was to disturb PSD-95 binding to Kal7 with interfering peptides. However, no competition could be demonstrated, despite various experimental approaches that included multiple resins and a wide range of concentrations of competing peptides. Covalent binding of PDZ123 to Affigel and to UltraLink Biosupport resins showed a dose-dependent saturable binding of Kal7 to the PDZ123 resin and little background binding to the BSA resin, using 5 pmol purified Kal7 input (Miller, Yan, Machida, et al., 2017) (**Fig**.**7A**, asterisks). However, competing GluN2B and Kal7 peptides had no effect on the binding of baculovirus Kal7 to the PDZ123 resin, even though binding of Kal7 to the BSA resin was minimal (**Fig**.**7B**).

These results suggested that the original hypothesis was incorrect and perhaps PSD-95 was interacting with Kal7 and GluN2B through additional sites other than the COOH-terminal PDZ binding domains. It is possible that multiple PDZ domains were in play, that interfering with a single domain was inadequate to disrupt the overall interaction, or that immobilizing either of the binding partners was too restrictive for specific interactions to occur (outlined in **Fig**.**7C**). We took advantage of the fact that Kal8 lacks the C-terminus PDZ binding domain of Kal7 but contains the rest of the Kal7 structure (**Fig**.**7D**), in order to address whether Kal7 and BluN2B could interact through additional sites besides the terminal PDZ binding domains. We utilized expression vectors for Kal7, Kal8 and PSD-95, separately transfected these into HEK293 cells, and made protein extracts. We then used co-immunoprecipitation to test for binding of solubilized PSD-95 with either gently solubilized Kal7 or Kal8 proteins (**Fig**.**7D**). The gels demonstrated that both Kal7 and Kal8 proteins interacted specifically with PSD95, regardless of whether or not they have a PDZ binding motif.

## Discussion

Cell-penetrating peptide mimetics of PDZ binding motifs have been shown to disrupt protein-protein interactions as diverse as connexins with zona occludens, multiple binding proteins with AMPA receptors, and various functions of serotonin receptors (Flores, Li, Bennett, Nagy, & Pereda, 2008; Flores et al., 2012; Lee, Liu, Wang, & Sheng, 2002; Luscher et al., 1999; Luthi et al., 1999; Nishimune et al., 1998; Pichon et al., 2010). Additional experiments on spinal cord demonstrated that Kal7-derived peptides could obliterate the usual LTP and LTD seen in spinal cord preparations (Lu et al., 2015), and that cell-permeant Kal7-derived peptides disrupt the accumulation of Kal7 in dendritic spines of cultured hippocampal neurons (Ma, Wang, et al., 2008).

Taken together, our current data on hippocampal slices extend previous studies that identified Kal7 and the GluN2B subunit-containing NMDA receptors as essential postsynaptic components of synaptic plasticity in CA1 neurons of mouse hippocampus. Further, we have integrated synthetic peptides, whole cell patch clamp electrophysiology and binding site investigations to probe the functional interactions between Kal7, GluN2B and PSD95 and their roles in mediating both LTP and LTD in pyramidal neurons in CA1 of the hippocampus. The striking finding is that synthetic peptides can affect synaptic plasticity, both long-term potentiation and long-term depression, within minutes after patching the neuron, arguing that a lifelong deficit in Kal7 (i.e. the Kal7 knockout mouse) is not necessary for at least some of the deficits in plasticity seen with the appropriate patterns of stimulation of the Schaffer collaterals.

Disrupting Kal7/PSD95, GluN2B/PSD95 and GluN2B/Kal7 interactions with peptide mimics blocks 7 train theta burst stimulation (TBS)-induced long term potentiation in CA1 of mouse hippocampus, while leaving LTP produced by high frequency stimulation (HFS) untouched. Further, disruptions of Kal7/PSD95 and NR2B/PSD95 binding, but not disruption of NR2B/Kal7, also block low frequency stimulation (LFS)-induced long-term depression. A recent paper demonstrated, by creating transgenic knock-in mice bearing intramolecular swaps of the COOH-terminal regions of GluA1 and GluA2, that the COOH-terminal domain of GluA1 was sufficient to enable the expression of LTP in response to HFS, even when the GluA1 COOH-terminal was attached to the GluA2 ion channel-transmitter receptor domains (Zhou et al., 2018). Similarly, the COOH-terminal domain of GluA2 was sufficient to facilitate the expression of LTD in response to high frequency stimulation, even when attached to the GluA1 ion channel-transmitter receptor domains (Zhou et al., 2018). Our electrophysiological results demonstrate consistent differences between LTP in response to HFS vs. LTP in response to TBS, which is generally thought to mimic more closely the signaling patterns in the animal (Larson & Munkacsy, 2015; Larson et al., 1986; Vanderwolf, 1969).

Several studies used patch recording pipettes to deliver a small peptide to interfere with the interaction of NSF (N-ethylmaleimide-sensitive fusion protein) and AMPA receptors, thereby blocking the development of LTD at hippocampal synapses (Luscher et al., 1999; Luthi et al., 1999; Nishimune et al., 1998), and further work also established a key role for AP-2 (clathrin adaptor protein-2) in LTD, at a distinct binding site (Lee et al., 2002). More recently, cell-penetrating peptide mimetics of the PDZ domain of 5HT2a were employed to relieve hyperalgesia, enhancing the potency of selective serotonin reuptake inhibitors (Pichon et al., 2010). The goldfish Mauthner cell system provided the model system to show that the binding of zona occludens (ZO-1) protein to the PDZ domain of connexin Cx35 was crucial for the synaptic delivery of the connexin to the gap junction synapse (Flores et al., 2008; Flores et al., 2012), demonstrating that gap junctions are extremely dynamic, with half-lives on the order of a few hours. In each of these studies, intracellular delivery of a peptide that interfered with protein-protein interactions was shown to have a rapid effect on synaptic electrophysiology.

We have used intracellular delivery of interfering peptides, along with binding affinity studies, to probe the role of Kalirin-7 in synaptic plasticity. The results demonstrated that the direct interactions of both Kal7 and the GluN2B subunit with a PDZ domain, presumably the abundant PSD-95, are essential for two major forms of synaptic plasticity, TBS-induced LTP and LFS-induced LTD. Strikingly, HFS-induced LTP, as used in many studies (e.g. (Zhou et al., 2018)), was not blocked by these interfering peptides. The direct interaction of the PH domain of Kal7 with the juxtamembrane domain of GluN2B was also shown to be the most delicately poised interaction for maintaining TBS-induced LTP, but strikingly played no role in LTD.

Our data illustrate that appropriate protein domain interactions are crucial for proper signal complex formation and localization to confer correct function. While specific interactions between binding sites may be suggested at the electrophysiological level of a single cell, larger scale biochemical experiments elucidate that these molecules have more complex binding capabilities. The notion that small molecules are able to disrupt specific interactions of PDZ domains may still present potential therapeutic applications without triggering greater changes at the PSD (Houslay, 2009).

## Acknowledgments

This work was supported by NIH DK032948 (REM & BAE).

